# Consequences of ignoring variable and spatially-autocorrelated detection probability in spatial capture-recapture

**DOI:** 10.1101/2020.12.24.424360

**Authors:** Ehsan M. Moqanaki, Cyril Milleret, Mahdieh Tourani, Pierre Dupont, Richard Bischof

## Abstract

**Context:** Spatial capture-recapture (SCR) models are increasingly popular for analyzing wildlife monitoring data. SCR can account for spatial heterogeneity in detection that arises from individual space use (detection kernel), variation in the sampling process, and the distribution of individuals (density). However, unexplained and unmodeled spatial heterogeneity in detectability may remain due to cryptic factors, intrinsic and extrinsic to the study system.

**Objectives:** We identify how the magnitude and configuration of unmodeled, spatially variable detection probability influence SCR parameter estimates.

**Methods:** We simulated realistic SCR data with spatially variable and autocorrelated detection probability. We then fitted a single-session SCR model ignoring this variation to the simulated data and assessed the impact of model misspecification on inferences.

**Results:** Highly autocorrelated spatial heterogeneity in detection probability (Moran’s *I* = 0.85 - 0.96), modulated by the magnitude of that variation, can lead to pronounced negative bias (up to 75%), reduction in precision (249%), and decreasing coverage probability of the 95% credible intervals associated with abundance estimates to 0. Conversely, at low levels of spatial autocorrelation (median Moran’s *I* = 0), even severe unmodeled heterogeneity in detection probability did not lead to pronounced bias and only caused slight reductions in precision and coverage of abundance estimates.

**Conclusions:** Unknown and unmodeled variation in detection probability is liable to be the norm, rather than the exception, in SCR studies. We encourage practitioners to consider the impact that spatial autocorrelation in detectability has on their inferences and urge the development of SCR methods that can take structured unknown or partially unknown spatial variability in detection probability into account.

## Introduction

Imperfect detection is one of the primary challenges to the estimation of the size of wild populations. Regardless of the data collection method employed, rarely, if ever, are all individuals in a population detected. This challenge is amplified with the increasingly widespread application of non-invasive sampling methods for making landscape-level assessments, such as camera trapping and non-invasive DNA sampling, which often trade off local search effort or capture intensity for spatial coverage (grain vs. extent; Chandler and Hepinstall-Cymerman 2016; Steenweg et al. 2018). Capture-recapture and, more recently, spatial capture-recapture (SCR) models estimate and account for imperfect detection, thereby producing robust estimates of the focal ecological parameters (Chao 2001; Efford 2004; Lukacs and Burnham 2005; Borchers and Efford 2008; Royle et al. 2014). SCR has become particularly popular as it exploits the information contained in the spatial configuration of detections and non-detections across the study area to produce spatially-explicit estimates of abundance, i.e. density (Borchers and Efford 2008; Royle and Young 2008; Royle et al. 2018).

In most field studies, detection probability – the probability of detecting a species or individual when it is present – is not only imperfect but also variable. Detection probability can vary across individuals, time, and space (Gimenez et al. 2008; Kellner and Swihart 2014; Conn et al. 2017; Guélat and Kéry 2018). The implications and treatment of individual and temporal variability in detection probability have been extensively documented in the non-spatial capture-recapture literature (e.g. Chao 2001; Link 2003; Borchers et al. 2006; Gimenez et al. 2018a). SCR uses spatial information and generates spatially-explicit predictions; thus, it has the potential to accomplish the same for spatial variability in detection probability (Efford et al. 2013; Royle et al. 2014). Spatial variation in detection probability as a result of the declining probability of detection with increasing distance from an individual’s activity center (AC) is in fact exploited by SCR models to estimate the position of latent ACs (Borchers and Efford 2008; Royle and Young 2008). However, there are other sources of spatial heterogeneity in the probability of detection, some caused by the study itself, such as variable search effort, and others due to local intrinsic and extrinsic factors that influence the detectability of animals (Efford et al. 2013; Efford and Mowat 2014; Royle et al. 2014). Some sources of spatial heterogeneity in detection probability are known, such as when sampling effort is recorded during camera trapping and non-invasive DNA sampling (Royle et al. 2009; Efford et al. 2013). Others are suspected and can be modeled with spatial covariates, such as habitat proxies for vulnerability to detection (Bischof et al. 2017; Kendall et al. 2019). SCR models are well-suited for these situations of variable detectability, as studies are usually configured into discrete detection locations referred to as detectors (or traps), which are distributed across a study area. Detectors can in turn be associated with detector-specific, and thus spatially explicit covariates that affect detection probability (Efford et al. 2013; Royle et al. 2014).

Nonetheless, in many wildlife monitoring studies, spatial heterogeneity in detection probability remains partially unknown. Unaccounted environmental factors may impact exposure to detectors; for example, site-specific characteristics may affect visibility, or local climate can influence genotyping success rate of non-invasively collected DNA samples (Efford et al. 2013; Kendall et al. 2019). Survey effort may also vary across the study area unbeknownst to the investigator. For example, many large-scale monitoring programs combine structured sampling with unstructured data collection methods to increase the spatial extent and sampling intensity, and involve members of the public in the process (Thompson et al. 2012; Conn et al. 2017; Altwegg and Nichols 2019; Bischof et al. 2020a; Sicacha-Parada et al. 2020). Data from unstructured sources introduce unknown spatial heterogeneity in detection probability in SCR studies. In the analysis of monitoring data, ignoring the variability in detection can seriously degrade population inferences (Nichols and Williams 2006; Gimenez et al. 2008; Gerber and Parmenter 2015). However, it has also been shown previously that spatially random variation in detection probability does not seem to be a major source of bias in parameter estimates from SCR analysis (Bischof et al. 2017).

Spatial autocorrelation in detection probability – when detectability is more similar among neighboring than distant detectors – is common in ecological studies (Guélat and Kéry 2018). Observed and unobserved spatially-autocorrelated variation in detection probability could have many causes (Gaspard et al. 2019), divided into two broad categories. On the one hand is the nature of the data collection; for example, regional differences in the mobilization of volunteers for non-invasive DNA collections, variation in camera trap efficiency due to inadvertent scent contamination at a cluster of sites, or reduced physical capture success in traps installed by a less-experienced operator in their designated area (Kristensen and Kovach 2018; Bischof et al. 2020a; Tourani et al. 2020a). On the other hand are characteristics of the study species and its environment, such as spatial gradients or clusters in site utilization (not density) or spatial variation in behavior (e.g. shyness) that lead to differential detectability (Efford et al. 2013; Howe et al. 2013; Lamb et al. 2020). Variable detection probability that is highly autocorrelated may disproportionately affect the overall detectability of certain individuals in the population based on the location of their ACs. In extreme cases, large swaths of the study area, and the individuals inhabiting them, may be left unsampled, unbeknownst to the investigator. We therefore expect flawed inferences from SCR studies, if such autocorrelated heterogeneity in detection probability remains cryptic, and thus unaccounted for. Specifically, when spatial autocorrelation in detection probability is not correlated with animal density (Clark 2019; Paterson et al. 2019), we predict that an SCR model assuming homogeneous detection probability would produce positively biased estimates of average detection probability and, therefore, negatively biased estimates of density.

Most, if not all, SCR analyses of empirical data ignore some sources of spatial variation in detection probability. The extent to which the misspecification of the detection process in the presence of variable and spatially-autocorrelated detection probability may affect the estimates of focal parameters in SCR studies has not yet been systematically investigated. Using simulations with an envelope that includes scenarios encountered in real-life wildlife monitoring, we quantify how unmodeled, spatially variable detection probability influences parameter estimates obtained via SCR. We do so with a focus on both the magnitude of variability in detection probability and the autocorrelation therein.

## Methods

### Spatial capture-recapture model

For the purpose of this study, we used a standard, single-session, Bayesian SCR model (Royle et al. 2014). Our model is composed of two hierarchical levels:

#### 1. The ecological sub-model

reflects the underlying ecological process of interest and describes the distribution of individuals in space, i.e. density. Following a homogeneous point process (Royle et al. 2014), we assume every individual in the population has a fixed AC (or home range center; *s*_*i*_) and that these individual ACs are randomly distributed across the habitat *S* according to a uniform distribution:

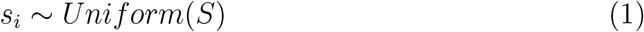

We used a data augmentation approach to account for those individuals in the population that are not detected (Royle et al. 2007). Detected and augmented individuals make up the super-population of size *M*. A latent state variable *z*_*i*_ describes inclusion of individual *i* in the population, governed by the inclusion probability *ψ*:

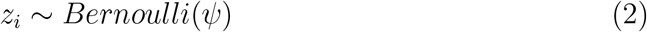

where *z*_*i*_ takes value 1 if individual *i* is a member of the true population and 0 otherwise. Population size *N* is therefore:

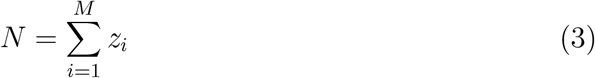

#### 2. The observation sub-model

describes the individual and detector-specific probability of detection *p*_*ij*_ as a function of euclidean distance *d*_*ij*_ between the location of an individual AC and a given detector *j*. We used the half-normal detection function to model *p*_*ij*_ (Borchers and Efford 2008; Royle et al. 2014):

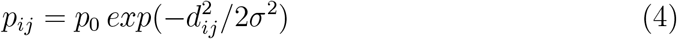

where *p*_0_ is the baseline detection probability (or magnitude of the detection function). In the half-normal model, the detection probability *p*_*ij*_ decreases monotonically with distance *d*_*ij*_ from *s*_*i*_ (Borchers and Efford 2008; Royle et al. 2014). The spatial scale parameter *σ* defines the rate of decline in *p*_*ij*_ with distance *d*_*ij*_ from detector *j* to the location *s*_*i*_ of AC. By including the spatial information through an explicit model for detection, SCR accounts for one important source of spatial variation in detection probability, the location of an individual’s AC relative to the detectors (Borchers and Efford 2008; Royle et al. 2014).

We considered the observations of individuals at detectors as the outcome of a Bernoulli process (detections [*y*_*ij*_ = 1] and non-detections [*y*_*ij*_ = 0]) with probability *p*_*ij*_ and conditional on the state *z*_*i*_ of individual *i*:

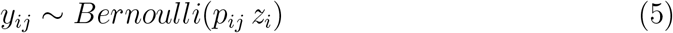

Importantly, this observation model assumes constant baseline detection probability among detectors and, thus does not account for additional detector-specific variation in detectability.

### Simulation

#### General approach

To evaluate the consequences of unmodeled spatial heterogeneity in detectability, we generated SCR data sets with varying patterns of spatial heterogeneity in the baseline detection probability (i.e. detector-specific *p*_0_), before fitting SCR models assuming constant baseline detection probability across detectors. We considered two main groups of scenarios: continuous and categorical variation in detectability. For each scenario, we varied both the level of spatial autocorrelation and the magnitude of the spatial variation to resemble sampling configurations and intensities that may occur in real-life studies (Fig. 1). We also included a baseline scenario without spatial heterogeneity in detection probability (thus, the observation sub-model was not misspecified) to use for reference.

**Figure 1:**
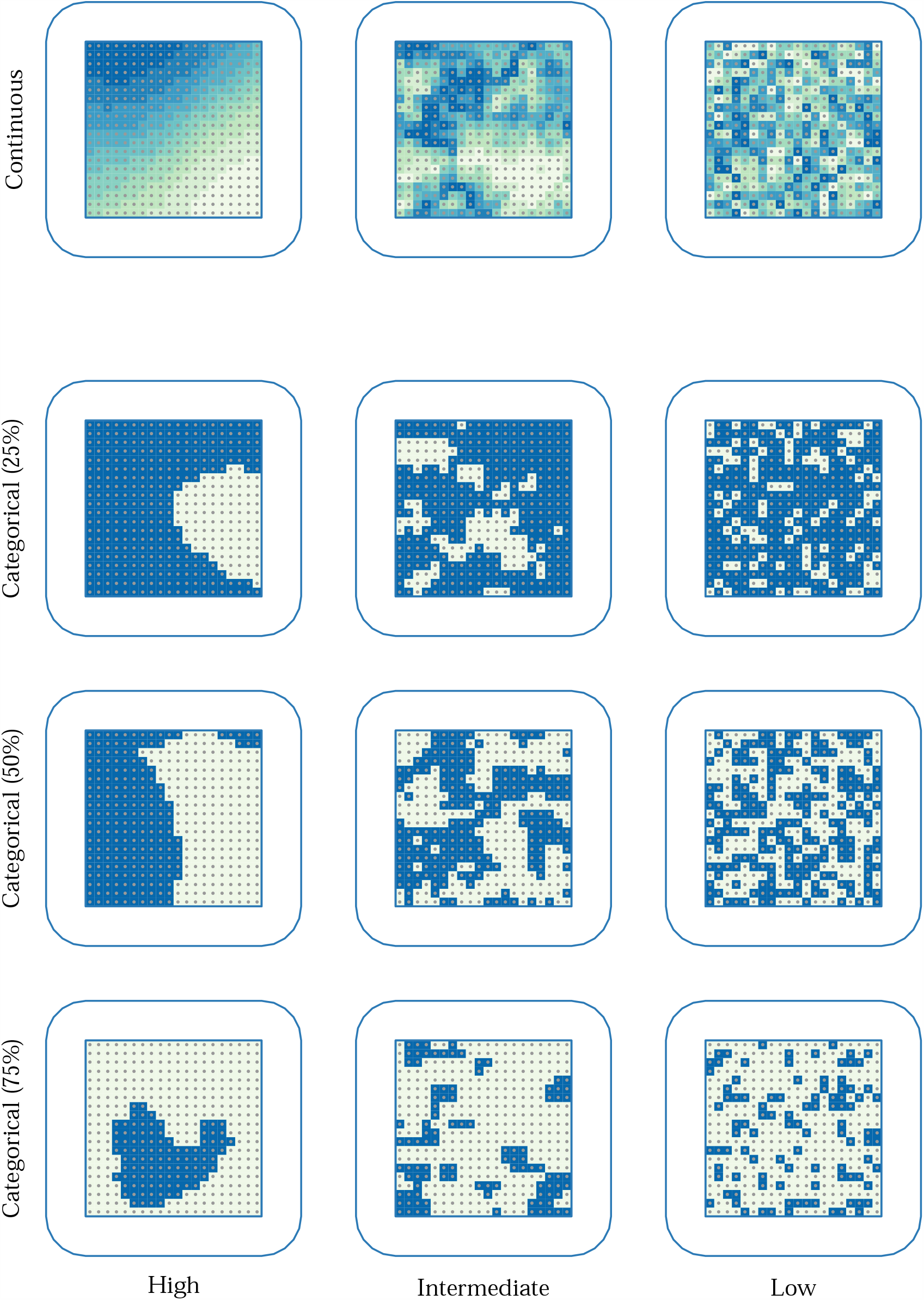
Examples of simulated, spatially variable and autocorrelated detection probabilities. Detectors (gray dots) are configured into a regular grid (inner polygon outline) centered on the habitat and surrounded by an unsearched habitat buffer shown as a white region between the inner and outer polygon outlines. Scenarios include continuous (top row) and discrete categorical variation (bottom three rows) in baseline detection probability, with lighter shading indicating lower baseline detection probability. Scenarios with continuous variation in baseline detection probability differ only in terms of the spatial autocorrelation in detection probability (decreasing from left to right). Scenarios with discrete categorical variation in baseline detection probability vary both in terms of proportions of detectors that belong to the group of detectors with lower detectability (increasing from top to bottom) and in terms of the spatial autocorrelation in detection probability (decreasing from left to right). High spatial autocorrelation illustrates a structured spatial covariate with values close to +1 (perfectly positively correlated), while Low indicates a random covariate with Moran’s *I* close to 0 (no autocorrelation).

#### Set up

For all simulations, we used a 20 × 20-distance unit (du) square grid of 400 detectors with 1 du inter-detector spacing. The habitat *S* included the region covered by detectors and a 4.5-du wide buffer around it (i.e. three times the simulated value for *σ*; Efford 2011) for a total area of 29 × 29 du^2^ (Fig. 1). The buffer allows individuals with ACs located outside but near the detector area to be detected within. We fixed the values for the true population size to *N* = 250 and the spatial scale parameter of the half-normal detection function to *σ* = 1.5 du across all simulation scenarios for simplicity. We set the size of the augmented population size to be 2.5 times the simulated number of ACs (*M* = 625).

#### Detector-level covariates

We generated spatially-autocorrelated covariates of the baseline detection probability *p*_0_ encompassing the extent of the 20 × 20 du^2^ detector grid with the same spatial resolution (1 du) using a function developed by Guélat (2013) with minor modifications. The covariate **X** was generated using a multivariate normal distribution:

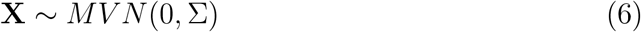

where the covariance matrix Σ determines the spatial association between detectors. Σ is calculated using a function *D* representing the decay in correlation between pairs of detectors with distance. We followed Guélat (2013) and modeled the covariance of **X** at two detectors *j* and *j*^*t*′^ as an exponential decay with distance:

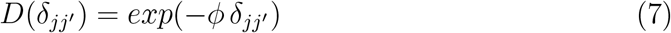

where *δ*_*jj*′_ is the distance between detectors *j* and *j*′, and *ϕ* is the rate determining how rapidly correlation declines with distance. We varied *ϕ* to simulate covariates with low (*ϕ* = 1000), intermediate (*ϕ* = 1), or high (*ϕ* = 0.001) spatial autocorrelation (Fig. 1). We randomly generated 100 covariate surfaces for each simulation scenario and scaled the resulting values. We then extracted the spatial covariate values for each detector grid cell **X**_*j*_ (but see below the extra step for simulating continuous variation in detectability). We further quantified the realized spatial autocorrelation using the Moran’s index of global spatial autocorrelation, Moran’s *I* (Moran 1950; Sokal and Oden 1978; Lichstein et al. 2002) for each simulated spatial covariate with the package ‘raster’ (Hijmans 2019). *I* ranges from −1 (perfectly negatively correlated) to 0 (no correlation) and 1 (perfectly positively correlated).

#### Simulation scenarios

##### 1. Continuous, detector-level variation in detectability

Continuous spatial variation in detection probability may arise in situations where an underlying habitat covariate, such as elevation, forest cover, or distance from a feature (e.g. roads or human settlements), linearly affects baseline detection probability at detectors (Fig. 1) but remains unmodeled. We modeled the detector-specific baseline detection probability 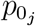 as a linear function of the simulated spatial covariate **X**:

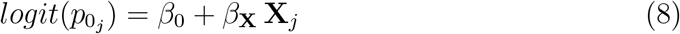

where *β*_0_ is the intercept value of the baseline detection probability *p*_0_ on the logit scale and *β*_**X**_ is the regression coefficient of the covariate effect. We kept *β*_0_ constant across simulations, which corresponds to an intercept value of 0.15 for *p*_0_. We used two values of *β*_**X**_ to generate low (*β*_**X**_ = −0.5) or high (*β*_**X**_ = −2.0) amounts of spatial variation in detection probability. Generating spatial covariates directly as spatially autocorrelated rasters as described above, leads to less tractable outcomes as the level of spatial autocorrelation *ϕ* influences the density distribution (not only the spatial distribution) of the covariate. To ensure that the density distribution of spatial covariates on detection probability remained comparable across simulations, regardless of the level of autocorrelation, we mapped a uniformly distributed spatial covariate with values in the range between −1.96 and 1.96 onto the spatially-autocorrelated similarity raster created as described above (see Appendix 1). Following Equation 8, the detector-specific baseline detection probability 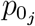 was therefore between 0.06 and 0.32 (median = 0.15) when the variation in detectability was low (*β*_**X**_ = −0.5). By increasing *β*_**X**_ to −2.0, 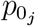 was between 0.003 and 0.9 (median = 0.15). *p*_*ij*_ is then calculated based on Equation 4:

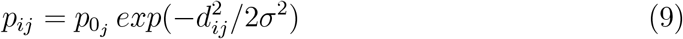

Finally, the SCR data *y*_*ij*_ is generated by realizing the detection process following Equation 5.

##### 2. Categorical, detector-level variation in detectability

In real-world monitoring studies, this situation arises when there are at least two regions with different sampling intensity across the study area; for example, two regions with varying sampling effort (or different sampling protocols that lead to varying sampling intensity), or two contrasting landscapes (e.g. forest vs. grassland) or detector types (e.g. camera traps on and off trails) across the study area that influence the detection probability (Fig. 1). To represent this situation, we transformed the underlying continuous spatial covariate into a binary one (**X**_*j*_ = 0 or 1) before simulating scenarios with two classes of detector-specific baseline detection probability *p*_0_ using Equation 8. The R code in Appendix 1 shows the method used to define the cut-off value to discretize the spatial covariates. We used the same values of *β*_**X**_ to generate low or high amount of variation in detectability between the discrete classes of detectors. With this set-up, the baseline detection probability at a group of detectors (**X**_*j*_ = 0) was equal to the intercept value 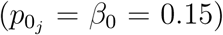, and 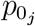 for the remaining detectors with lower detectability (**X**_*j*_ = 1) was either 0.1 (when *β*_**X**_ = −0.5) or 0.02 (*β*_**X**_ = −2.0) depending on the simulated amount of spatial variation in detectability. In addition, we considered an extreme case, where a portion of the study area remained entirely unsampled. This situation could be encountered in, for example, volunteer-based monitoring programs, where SCR data are collected opportunistically but no or only limited spatial information exists about the spatial configuration of sampling. Logistic issues, such as systematic equipment failure (e.g. vandalism) or human error, may also result in clusters of detectors remaining unsearched or inactive. We set *β*_**X**_ to −10000, so that 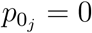 for detectors with a covariate value of **X**_*j*_ = 1, and therefore no detection could occur at these inactive detectors.

To evaluate the potential effect of the proportion of detectors with a different baseline detection probability, we varied the proportions of detectors assigned to each class of the discrete covariate simulated (Fig. 1). We simulated SCR data sets with 25% (n = 100 detectors), 50% (n = 200), or 75% (n = 300) of the detectors assigned to the lower detectability class (**X**_*j*_ = 1).

#### Simulation realization

In total, we generated six simulation scenarios of continuous variation in baseline detection probability with the combinations of two values of *β*_**X**_ and three levels of spatial autocorrelation *ϕ*. For the discrete categorical variation in baseline detection probability, we simulated 27 scenarios from all possible combinations of the three values of *β*_**X**_, three different proportions of detectors with lower detectability, and three levels of spatial autocorrelation. Finally, we included a scenario of constant detection probability across detectors for reference, i.e. the observation sub-model was correctly specified. We repeated the data simulation procedure 100 times for each combination of parameters, resulting in a total of 3400 simulated SCR data sets (Table S1, Appendix 2).

#### SCR model fitting and evaluation of model performance

We fitted the SCR model described earlier, which does not account for spatial variability in baseline detection probability, to the simulated data sets using NIMBLE (version 0.8.0; de Valpine et al. 2017) and R 3.6.1 (R Core Team 2019). We used a local evaluation approach to reduce computation time (Milleret et al. 2019). We drew from 3 chains, 15000 Markov chain Monte Carlo (MCMC) samples each, and discarded the initial 5000 samples as burn-in. We visually inspected the mixing of the chains using trace-plots and considered models as converged when all parameters had a potential scale reduction value (Rhat) *<* 1.10 (Brooks and Gelman 1998). We removed from further analysis simulation runs that had failed to reach convergence. R code exemplifying the SCR data simulation for each scenario and model fitting are provided in Appendix 1.

To quantify the consequences of unaccounted spatial heterogeneity in detection probability for SCR models, we calculated the relative bias (RB), coefficient of variation (CV), and coverage probability of the 95% credible intervals (hereafter, coverage; Walther and Moore 2005) of the estimates of population size 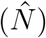 and spatial scale parameter 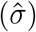. RB was calculated as:

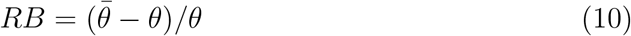

where 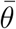 is the mean estimate for the parameter of interest and *θ* is the true (simulated) value of that parameter. CV was calculated to assess the precision of each parameter estimate (*θ*) as:

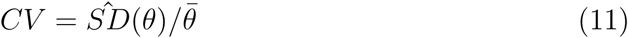

where 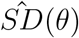 is the standard deviation of the estimate for that parameter (Walther and Moore 2005). Further, we calculated the coverage as the proportion of simulation runs in which the 95% credible interval of the parameter estimate included the simulated value of that parameter (Walther and Moore 2005).

## Results

### Overview

Of the 3400 simulation runs, 3384 (99.5%) reached convergence and were retained: 599 (99.8%) and 2685 (99.4%) for the continuous and categorical scenarios of variation in detectability, respectively (Table S2, Appendix 2). Our results indicate that, regardless of the simulation scenario, population size 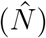 and spatial scale parameter 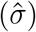 estimators were exhibited pronounced bias in the presence of high spatial autocorrelation among detectors (Fig. 2). Precision (Fig. 3) and coverage (Fig. 4) were also affected when the detector-specific variation in detection probability remained unmodeled. The consequences of spatial autocorrelation were amplified with increasing proportions of detectors with lower detectability (the categorical scenarios) and, in certain situations, when the amount of variation in detection probability was high (Table S2, Appendix 2).

**Figure 2:**
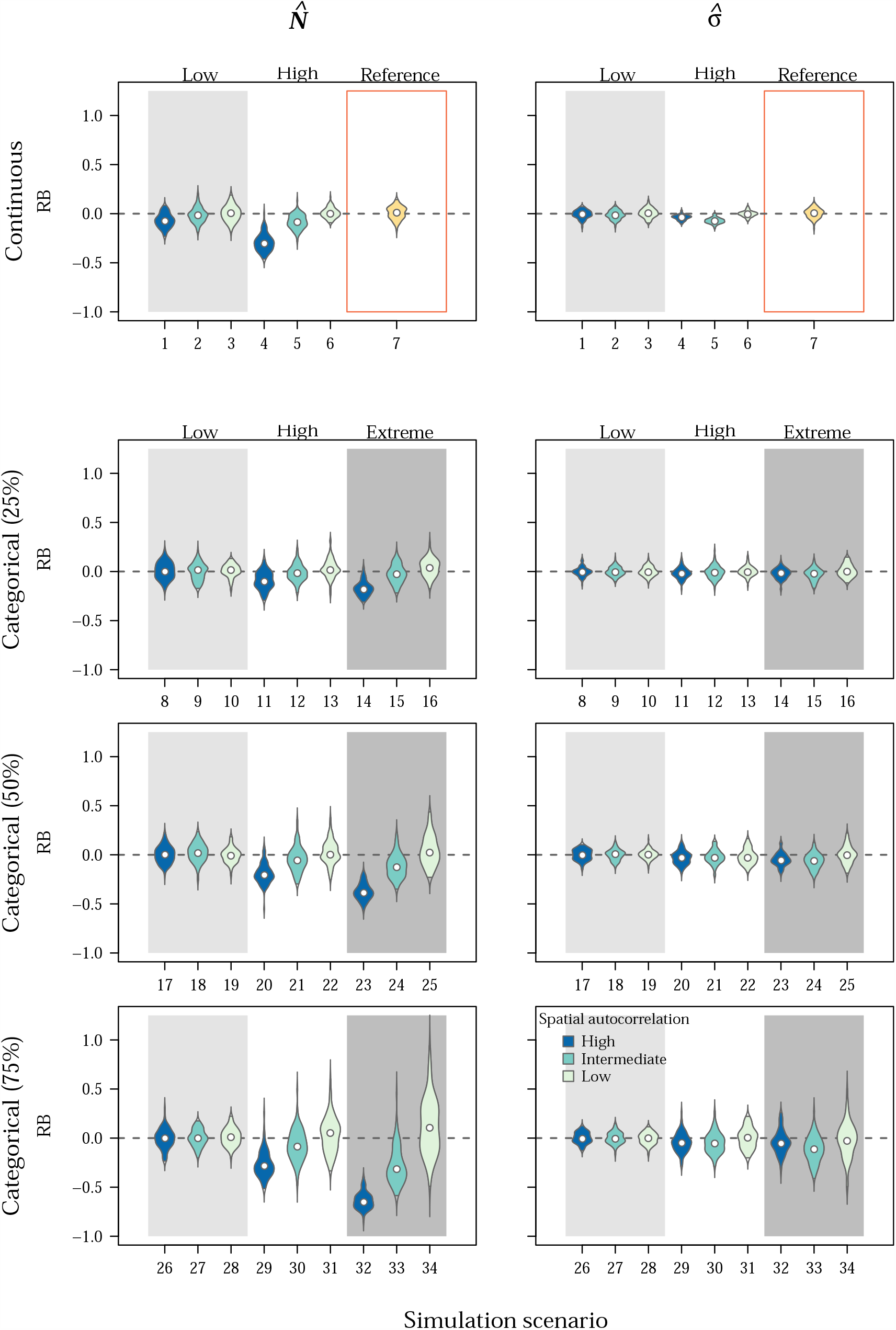
Relative bias (RB) for population size 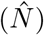 and the spatial scale parameter of the half-normal detection function 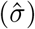 estimated by a spatial capture-recapture model fitted to simulated data sets. Spatial variation in detection was simulated but not accounted for in the estimation model. Violins show the biasing effects of spatial autocorrelation at the detector-level covariates simulated (decreasing from dark [High] to light colors [Low]) in different scenarios of continuous (top row) and discrete categorical (bottom three rows) variation in baseline detection probability. For the categorical scenarios, the proportions of detectors that belong to a group of detectors with lower detectability increases from top (25%) to bottom (75%). Background colors correspond to three different sets of simulation scenarios with similar values of the magnitude of the covariate effect: Low (*β*_**X**_ = − 0.5; light gray), High (*β*_**X**_ = − 2.0; white), and Extreme (*β*_**X**_ = − 10000; dark gray). The yellow violins labeled as Reference in the top row show the results for a baseline scenario without heterogeneity in detection probability.

**Figure 3:**
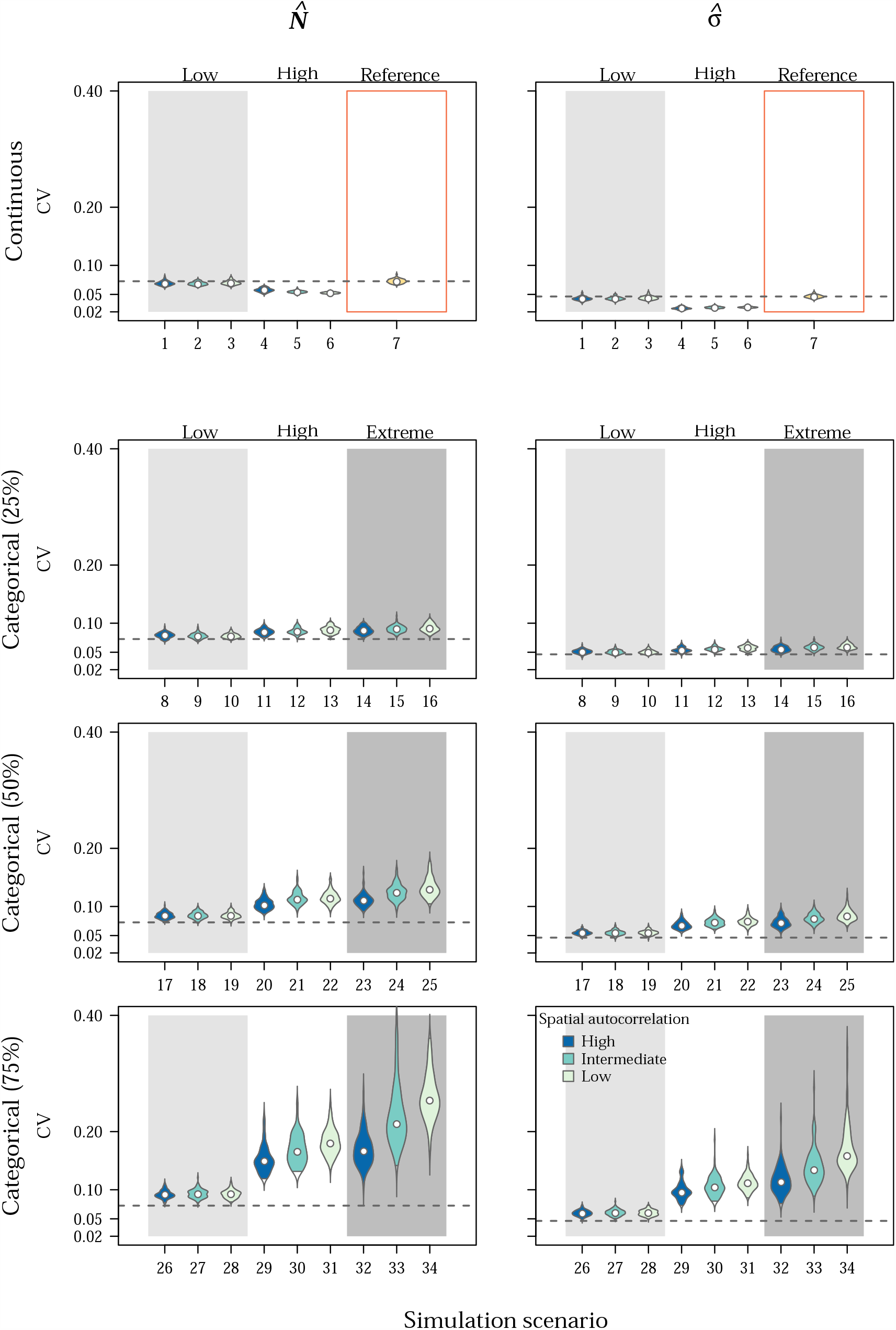
Coefficient of variation (CV) for population size 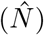 and the spatial scale parameter of the half-normal detection function 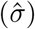 estimated by a spatial capture-recapture model fitted to simulated data sets. Spatial variation in detection was simulated but not accounted for in the estimation model. Violins show the effects of spatial autocorrelation in the detector-level covariates simulated (decreasing from dark [High] to light colors [Low]) in different scenarios of continuous (top row) and discrete categorical (bottom three rows) variation in baseline detection probability. For the categorical scenarios, the proportions of detectors that belong to a group of detectors with lower detectability increases from top (25%) to bottom (75%). Background colors correspond to three different sets of simulation scenarios with similar values of the magnitude of the covariate effect: Low (*β*_**X**_ = − 0.5; light gray), High (*β*_**X**_ = − 2.0; white), and Extreme (*β*_**X**_ = − 10000; dark gray). The yellow violins labaled as Reference in the top row show the results for a baseline scenario without heterogeneity in detection probability, which was included for reference. The dashed lines show median CV for the parameter estimates achieved in the reference scenario.

**Figure 4:**
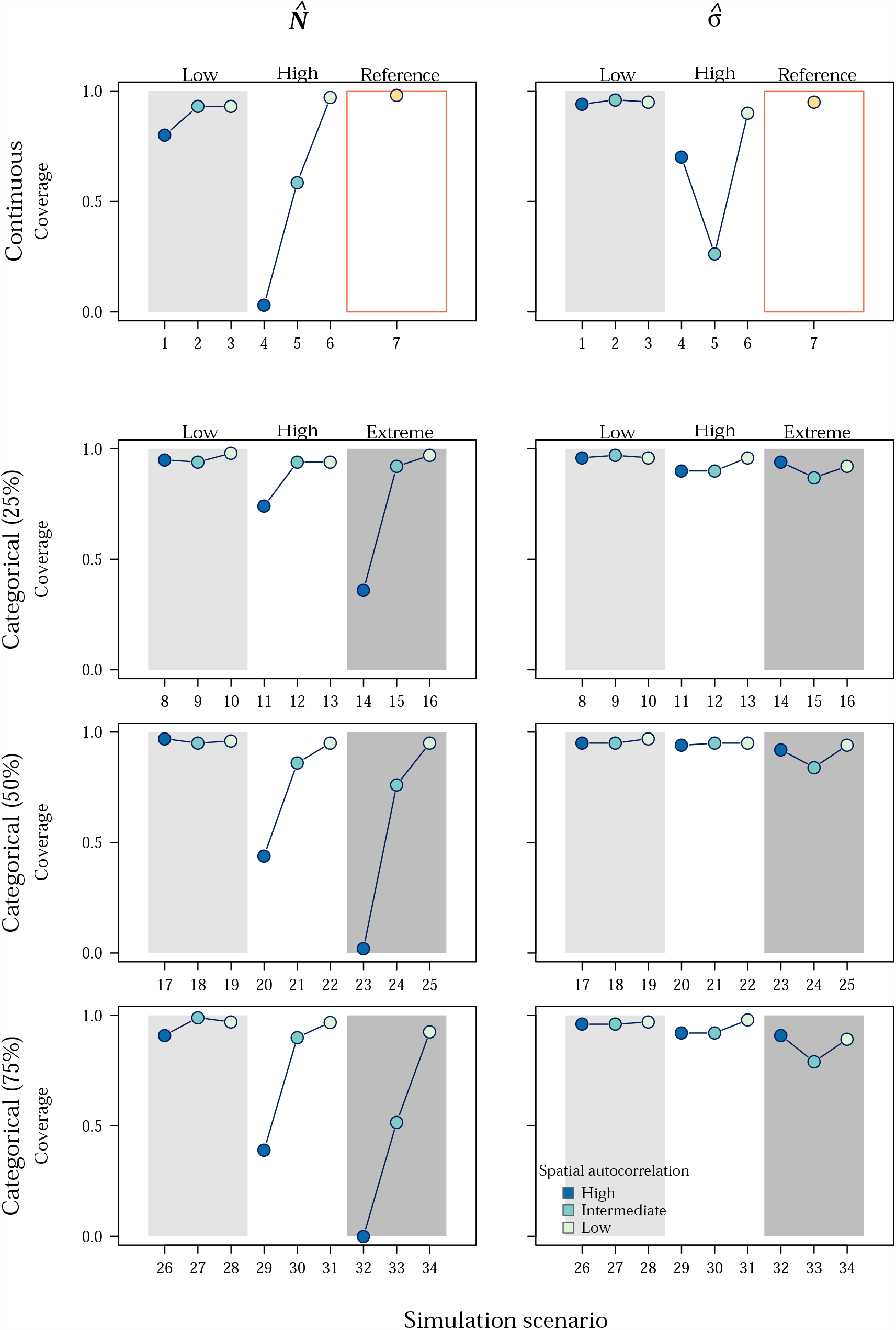
Coverage probability of the 95% credible intervals of population size 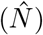 and the spatial scale parameter of the half-normal detection function 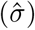 estimated by a spatial capture-recapture model fitted to simulated data sets. Spatial variation in detection was simulated but not accounted for in the estimation model. Points show the effects of spatial autocorrelation of continuous (top row) or discrete categorical (bottom three rows) baseline detection probabilities (decreasing from dark [High] to light colors [Low]). For the categorical scenarios, the proportions of detectors that belong to a group of detectors with lower detectability increases from top (25%) to bottom (75%). Background colors correspond to three different levels of variation in the baseline detection probability: Low (*β*_**X**_ = − 0.5; light gray), High (*β*_**X**_ = − 2.0; white), and Extreme (*β*_**X**_ = − 10000; dark gray). The yellow points labeled as Reference in the top row show results for a baseline scenario without heterogeneity in detection probability.

### Continuous, detector-level variation in detectability

We observed increasingly negative bias in 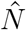 with increasing spatial autocorrelation (Fig. 2; Table S2, Appendix 2). The magnitude of RB in 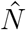 obtained with high spatial autocorrelation (simulation scenarios 1 and 4: median Moran’s *I* = 0.96; median 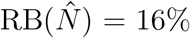) was substantially greater than that for scenarios with low spatial autocorrelation (scenarios 3 and 6: median Moran’s *I* = 0; median 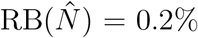). This pattern was amplified when the amount of variation in detectability was high (scenario 4: median 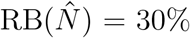). In contrast, we detected no noticeable bias in 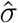 (Fig. 2), even when the amount of variation in detectability was high and spatial autocorrelation was high or intermediate (scenarios 4 and 5: median 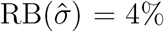 and 7%, respectively).

The pattern in precision of 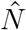 and 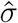 were almost identical across the scenarios considered (Fig. 3). When the amount of variation in detectability was low (scenarios 1-3), precision of the parameter estimates were comparable to those of the reference scenario without spatial heterogeneity in detection probability (scenario 7: median 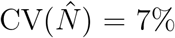 and median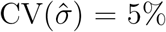). Precision of 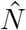 and 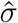 were inflated when the magnitude of variation in detectability was high (scenarios 4-6: median 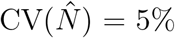 and median 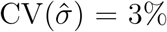), where increasing the spatial autocorrelation slightly decreased the precision of 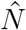). (scenario 4; Table S2, Appendix

Coverage of both 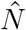 and, to a lesser extent, 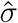 were drastically impacted by spatial autocorrelation (Fig. 4; Table S2, Appendix 2). In situations of high spatial autocorrelation and low variability in detection probability, we observed a 14% reduction in coverage of 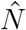, compared to low spatial autocorrelation (from Coverage 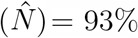 in scenario 3 to Coverage 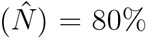 in scenario 1). The combination of high spatial autocorrelation and high variation in detectability led to coverage of 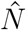 as low as 3% (scenario 4; Fig. 4). The pattern was less pronounced for 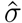 (scenario 4: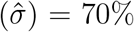). However, we detected a drastic decline in the coverage of 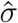 for the scenario of intermediate spatial autocorrelation and high amount of variation in detectability (scenario 5: median Moran’s *I* = 0.63,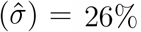; Fig. 4).

### Categorical, detector-level variation in detectability

We observed increasing negative bias in 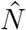 with increasing spatial autocorrelation and increasing proportion of detectors with lower detectability (Fig. 2; Table S2, Appendix 2). The bias was particularly pronounced in scenarios of extreme, spatially-autocorrelated variation in detectability (median Moran’s *I* = 0.86), where 50% or more of detectors were inactive, ultimately leading to an entire region with little sampling (scenarios 23 and 32: median 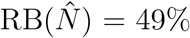). However, when the variation in detectability was low (scenarios 8-10, 17-19, and 26-28), 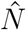 was minimally affected in terms of bias regardless of the spatial autocorrelation and proportions of detectors with lower detectability (Table S2, Appendix 2). Similarly, when spatial autocorrelation in detectability was low, 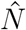 remained relatively unbiased even in the extreme scenarios of a portion of detectors being inactive. We observed no systematic bias in 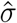 under any of the scenarios tested (Fig. 2; Table S2, Appendix 2).

Precision of 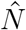 and 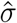 decreased with increasing proportions of detectors with lower detectability and increasing variation in detectability (Fig. 3; Table S2, Appendix 2). However, precision increased with spatial autocorrelation. For the extreme scenario of a partially-sampled area, when 75% of the detectors were inactive and spatial autocorrelation was low (scenario 34; Fig. 3), median 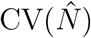 and median 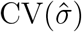 increased by 249% and 242%, respectively, compared to the reference scenario. In contrast, the increase in median 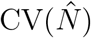 and median 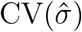 was 129% and 145%, respectively, for the scenario of high spatial autocorrelation (scenario 32; Table S2, Appendix 2).

The coverage of 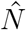 drastically decreased with increasing spatial autocorrelation (Fig. 4). With high spatial autocorrelation, the coverage of 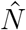 dropped to between 39% and 44% in the presence of high variation in detectability and higher proportions of detectors with lower detectability (scenarios 29 and 20). When more than 50% of the detectors were inactive, coverage of 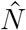 was near 0 (scenarios 23 and 32; Fig. 4). Coverage was nominal for 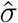 under low and high levels of spatial autocorrelation, and only decreased to between 79% and 87% when spatial autocorrelation was intermediate (scenarios 15, 24, and 33: median Moran’s *I* = 0.42; Table S2, Appendix 2).

## Discussion

Our study revealed that variable detection probability, which is ubiquitous in wildlife monitoring studies, can have pronounced consequences for inferences from SCR analyses. The critical factor is spatial autocorrelation in detection probability: highly autocorrelated and variable detectability leads to pronounced bias and reduction in precision and coverage of SCR estimates of population size, the main parameter of interest in such studies (Royle et al. 2014, 2018). Conversely, at low levels of spatial autocorrelation, SCR model estimates remained robust even to high amounts of unmodeled heterogeneity in detection probability. Estimates of the spatial scale parameter of the half-normal detection function were, however, unbiased and coverage remained comparatively high for most scenarios of spatially-autocorrelated detection probability. Nonetheless, the pattern of reduction in precision of the estimates of the scale parameter was similar to those in the estimates of population size.

SCR is a powerful analytical tool for spatially-explicit inference on the ecology of wild populations while accounting for imperfect detection (Royle et al. 2018). One of the advantages of SCR models over non-spatial capture-recapture models is the ability to address the effects of known spatio-temporal variation in detectability as a result of the spatial configuration of ACs relative to detectors and detectorspecific characteristics; thus, providing a more realistic model of the detection process (Efford et al. 2013; Royle et al. 2014). Yet, despite the spatially-explicit nature of SCR data collection and analysis, it is liable to be afflicted by unknown or undocumented variation in detection probability; for example, due to inadvertent (e.g. equipment failure, within-individual variations) or design-induced factors (e.g. non-random or preferential sampling), or as the result of data aggregation across multiple survey methods or detectors (Efford et al. 2013; Howe et al. 2013; Efford and Mowat 2014; Royle et al. 2014; Gerber and Parmenter 2015). As we have shown here, failure to adequately account for variable and spatially-autocorrelated detection probability may bias SCR parameter estimates, impact estimates of precision, and generally lead to erroneous inferences. In extreme cases among the ones we considered, this led to up to a 75% bias and to virtually zero probability that the credible interval contains the true value of population size (Table S2, Appendix 2). A misspecified detection model may therefore not only impact inferences, but also reduce the chance of meeting the management and conservation objectives that often motivate these studies.

We observed the strongest systematic bias in cases where a substantial and distinct region of the detector space was unsampled, essentially leaving a hole in the detector grid that remained unaccounted for in the model. The pronounced negative bias in 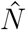 that arises in these situations can be explained if we consider individuals with ACs within such a hole have a high chance of being missed entirely during sampling, essentially creating a latent class of individuals with very low or zero detectability. As a consequence, detection probability estimates are biased high as they are based primarily on recaptures of detected individuals, and estimates of abundance and density are consequently biased low. This pattern is amplified by the magnitude of variation in detection probability between regions of lower and higher detectability, and by the size of the region with lower detection probability (Table S2, Appendix 2).

Situations where entire regions of the study area remain unsampled are fairly common in monitoring studies, where the data is gathered opportunistically, either in part or as a whole (Conn et al. 2017; Altwegg and Nichols 2019; Sicacha-Parada et al. 2020). Unstructured sampling can augment information, and thus improve population inferences about rare or elusive species (Thompson et al. 2012; Tenan et al. 2017; Sun et al. 2019; Bischof et al. 2020a). Opportunistically-collected data obtained as part of public surveys (e.g. citizen-science data) are sometimes integrated into monitoring programs as this allows investigators to sample areas at unprecedented scales with lower costs and the added benefit of public involvement in management and conservation practices (Altwegg and Nichols 2019; Bischof et al. 2020a; Sicacha-Parada et al. 2020). To minimize and account for variation in detection probability in such sampling schemes, volunteers should be encouraged to visit all habitats within the study area, reduce variability in observer proficiency by providing standardized training, and collect data on potentially relevant covariates (Altwegg and Nichols 2019).

This study was motivated by our previous work on large-scale monitoring of large carnivores, where spatial variation in effort can be challenging to quantify because the data are collected in a quasi-systematic fashion by field personnel associated with multiple jurisdictions and opportunistically by volunteer members of the public (Bischof et al. 2016, 2017, 2020a; Milleret et al. 2020). Although we considered a wide range of scenarios in terms of the magnitude and spatial configuration of variability in detection probability, we expect the consequences of unmodeled heterogeneity to be relaxed when detection probability is relatively high. In such situations, unknown and unmodeled variation in detectability is likely to be less problematic as individuals from the population are more likely to be detected at multiple detectors (i.e. increase in both the proportion of detected individuals from the population and spatial recaptures). The effects of spatial heterogeneity in detectability may also be mediated by the species space-use characteristics. Here, we simulated a target population with intermediate home-range overlap (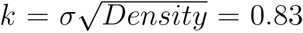; Efford et al. 2016) with a density of 0.3 AC/du^2^, which was constant across simulation scenarios. For species with small home-range sizes, where the spatial scale parameter is small relative to the distance between detectors, unmodeled spatially-autocorrelated detection probability may lead to more problematic inferences due to the increased risk of individuals going completely undetected in the resulting “holes” in the detector grid. In addition, we assumed homogeneous population density across the study area, which is rarely, if ever, the case in wild populations. Previous studies have suggested that the consequences of unmodeled heterogeneity in detection probability could be amplified when detectability is correlated with density (e.g. Clark 2019; Paterson et al. 2019). As a correlation between animal density and detection probability results in a positive bias in estimates of population size, we expect that the biasing effects we detected would be comparatively less pronounced when the unmodeled detection probability is spatially autocorrelated. However, the misspecification of the observation sub-model may lead to confounding effects between variation in detection probability and animal density that must be accounted for (Efford et al. 2013).

When the drivers of spatial variation in detection probability are known or suspected, the variation can be accounted for by, for example, including relevant covariates (e.g. transect length, camera trap nights, habitat or line-of-sight visibility) in the observation sub-model of SCR models (Royle et al. 2009, 2014; Efford et al. 2013; Efford and Mowat 2014; Bischof et al. 2017, 2020a). In the absence of proxies that can serve as fixed or random effects, how should investigators deal with unknown spatial heterogeneity in detectability in SCR analyses? One solution would be to discard affected data, such as observations contributed by members of the public without corresponding measures of search effort. This, however, would lead to a loss of potentially valuable information, both in terms of number of individuals detected and spatial recaptures (Marques et al. 2011; Sollmann et al. 2012; Tourani et al. 2020b). Explicitly modeling spatially-autocorrelated detectability, for example using conditional autoregressive (CAR) models, may offer a solution. CAR models have been implemented in other hierarchical modeling frameworks, such as non-spatial capture-recapture and spatial distribution modeling (e.g. Johnson et al. 2013; Chen and Ficetola 2019; Nicolau et al. 2020), but we are not aware of similar extensions in SCR studies. The added complexity resulting from the inclusion of an autoregressive component on a latent variable like detection probability could pose a significant computation barrier to implementation in Bayesian SCR at large spatial scales. However, recent advances in both software (de Valpine et al. 2017; Bischof et al. 2020b) and Bayesian SCR model formulations (Milleret et al. 2018, 2019; Turek et al. 2020) have improved model fitting efficiency, allowing for fitting SCR models of unprecedented complexity and spatial scales (Bischof et al. 2020a); thus, opening new possibilities.

Goodness-of-fit tests could help practitioners diagnose potential violations of model assumptions, including unmodeled spatial variation in detection probability and determine whether there is a need to account for it in the model in the first place. Such diagnostics should be an integral part of any ecological modeling exercise (Conn et al. 2018), and Bayesian p-values (Gelman et al. 1996) have been proposed as a general framework for goodness-of-fit testing of SCR models (Royle et al. 2014). However, goodness-of-fit diagnostics for SCR is a developing field of research and specific tests and recommendations, such as those that were developed for non-spatial capture-recapture models (e.g. Gimenez et al. 2018b) are still lacking. Thus, testing and accounting for the possible violations, as well as correcting the model estimates, are still challenging in SCR.

## Conclusions

Unmodeled spatially-autocorrelated variation in detection probability can noticeably impact the reliability of inferences derived using SCR. This specifically affects estimates of abundance and density, primary parameters in wildlife monitoring studies, and can therefore have severe consequences for wildlife management and conservation. Unobserved variability in detection probability is likely ubiquitous in real-life SCR studies, and we encourage research to develop approaches that help practitioners diagnose and account for it.

## Author contributions

Conceptualization: RB; Design and methodology: EM, RB, MT, CM, PD; Analysis: EM; Writing - original draft preparation: EM; Writing - review and editing: EM, RB, CM, MT, PD.

## Acknowledgements

This study received funding from the Norwegian Research Council grant 286886 (project WildMap). Thanks to Soumen Dey for helpful discussions.

## Supplementary Information

Supplementary material related to this article can be found, in the online version, at xx

